# A wireless, 60-channel, AI-enabled neurostimulation platform

**DOI:** 10.1101/2025.02.04.635298

**Authors:** Daniel S. Rizzuto, Haydn G. Herrema, Zhe Hu, Daniel Utin, Joshua Kahn, Chris Ho, Andrew Smiles, Robert E. Gross, Bradley C. Lega, Sandhitsu R. Das, Michael J. Kahana

**Affiliations:** Nia Therapeutics, Inc., Allston, 02134, MA, USA; Department of Neurosurgery, Rutgers University, New Brunswick, 08901, NJ, USA; Department of Neurology and Neurotherapeutics, University of Texas Southwestern, Dallas, 75390, TX, USA; Department of Neurology, University of Pennsylvania, Pennsylvania, 19104, PA, USA

**Keywords:** brain computer interface, neuromodulation, artificial intelligence, human memory, biomarkers

## Abstract

**Objective:** Closed-loop neuromodulatory therapies require devices that can both decode ongoing brain states and deliver multi-site stimulation.

**Methods:** We describe the Smart Neurostimulation System (SNS), a cranially mounted implant with 60 configurable recording/stimulation channels, inductive power, and onboard spectral-feature classification. In three freely-moving sheep we streamed local-field potentials and conducted two parameter sweep experiments.

**Results:** Cross-validated movement classifiers achieved an average AUC *>* 0.95. Increasing stimulation amplitude and frequency produced post-stimulation elevations in *α*-band (8-12 Hz) and *γ*-band (78-82 Hz) power at most target locations.

**Conclusion:** The SNS unifies high-density sensing, real-time brain state decoding, and programmable closed-loop stimulation in a single device, demonstrating behavioral-state prediction and parameter-dependent neuromodulation in vivo.

**Significance:** These findings establish a preclinical foundation for biomarker-guided stimulation targeting distributed cortical networks underlying memory and cognition.

**Highlights:**

- Wireless 60-channel implant enables simultaneous sensing and stimulation
- vine study demonstrates reliable classification of movement and stimulation-related modulation of spectral activity
- The SNS represents a platform for biomarker-guided neuromodulation therapies

## 1. Introduction

Closed-loop neuromodulation—which adapts in real time to the brain’s evolving state—offers a path to more effective and safer therapies across a wide range of neurological conditions. In epilepsy, responsive neurostimulation cuts seizure burden by stimulating the brain only when it senses preictal EEG patterns [1]. In Parkinson’s disease and essential tremor, adaptive deep-brain stimulation raises efficacy while reducing dyskinesias [2]. In chronic pain and depression, state-contingent stimulation promises symptom relief without the habituation and mood swings seen with fixed schedules [3, 4, 5, 6]. These gains arise because closed-loop devices tailor pulse timing, location, and amplitude to moment-by-moment neural activity rather than relying on clinician-programmed settings that may be optimal only under the static conditions present at the time of programming.

Despite these clinical gains, today’s adaptive systems still rely on a strikingly small information footprint. Medtronic’s *Percept* ^TM^ aDBS, for example, modulates amplitude from a single *β*-band power estimate per subthalamic lead, whereas the NeuroPace *RNS* ^®^ system for epilepsy responds to band-pass or line-length thresholds on up to four bipolar recording channels. Such low-dimensional control cannot capture the distributed, multifrequency dynamics that underlie higher-order functions such as memory, attention, or mood. Treating these complex indications, therefore, requires sampling many sites spanning distributed brain networks, extracting numerous spectral and temporal features simultaneously, and combining those features with a real-time classifier capable of predicting momentary fluctuations in neurocognitive state. Prior work has shown that multivariate classifiers trained on hundreds of spectral features can reliably decode moments of high versus low mnemonic efficacy in humans [7, 8, 9, 10]. Delivering such classifier-based closed-loop therapy, however, demands implants that record from many distributed sites, process spectral features onboard, and perform sensing and stimulation concurrently, all while remaining small, wireless, and power-efficient for chronic use.

Here we describe progress toward developing and testing a wireless, 60-channel, AI-enabled brain–computer interface, the Smart Neurostimulation System (SNS). Our proof-of-concept studies of closed-loop neuromodulation to improve memory motivated the SNS design [11, 12]. These studies, conducted in epilepsy patients with and without a prior history of traumatic brain injury (TBI), used external devices connected to a host PC to record brain activity and to apply stimulation through commercially available, FDA-cleared electrodes. Neurosurgeons implanted these electrodes semi-chronically for seizure and functional mapping to inform potential resective surgery.

Mnemonic ability varies not only from person to person but also from trial to trial within the same individual. Study of these fluctuations documented that known experimental variables accounted for only a small fraction of the stochastic variability, with the largest predictor being performance on the prior trial [13]. Classifiers trained on spectral features of intracranial EEG recordings predicted memory performance on both individual items and entire lists [14, 15]. These classifiers leveraged information recorded from widespread regions of the memory network, including lateral temporal, medial temporal, and prefrontal cortices.

Mnemonic effects of electrical stimulation depend on brain state: stimulation impaired memory when delivered during classifier-predicted good states and improved memory when delivered during classifier-predicted poor states [7]. In a subsequent study, we validated these observations by designing a closed-loop system that triggered high-frequency (100–200 Hz) stimulation bursts upon detected memory lapses, reliably improving memory for stimulated items [8]. A replication in a cohort (*N* = 8) of epilepsy patients with moderate-to-severe TBI further showed that the mnemonic benefits of closed-loop stimulation accrue to the entire stimulated list, not just to the stimulated items [11]. Analyses of a larger cohort of patients (*N* = 47) found that only stimulation near white-matter tracts yielded consistent list-level mnemonic benefits; among these targets, those with the strongest functional connectivity to the memory network produced greater mnemonic boosts [12]. These studies provide proof of concept for a brain–computer interface therapy for memory loss. Approved devices, however, cannot meet the multichannel sensing and closed-loop stimulation requirements of such a therapy. To satisfy these requirements, we designed the SNS.

## 2. The Smart Neurostimulation System (SNS)

Figure 1 illustrates the overall system design. The SNS comprises a cranially mounted implantable pulse generator (IPG) for processing neural data and controlling therapy delivery; a wearable external processor (EP) that powers the IPG and acts as a communication hub; four custom depth leads for sensing and stimulating neural tissue; and a physician programmer and cloud-based AI platform that personalizes patient therapy.

**Figure 1:**
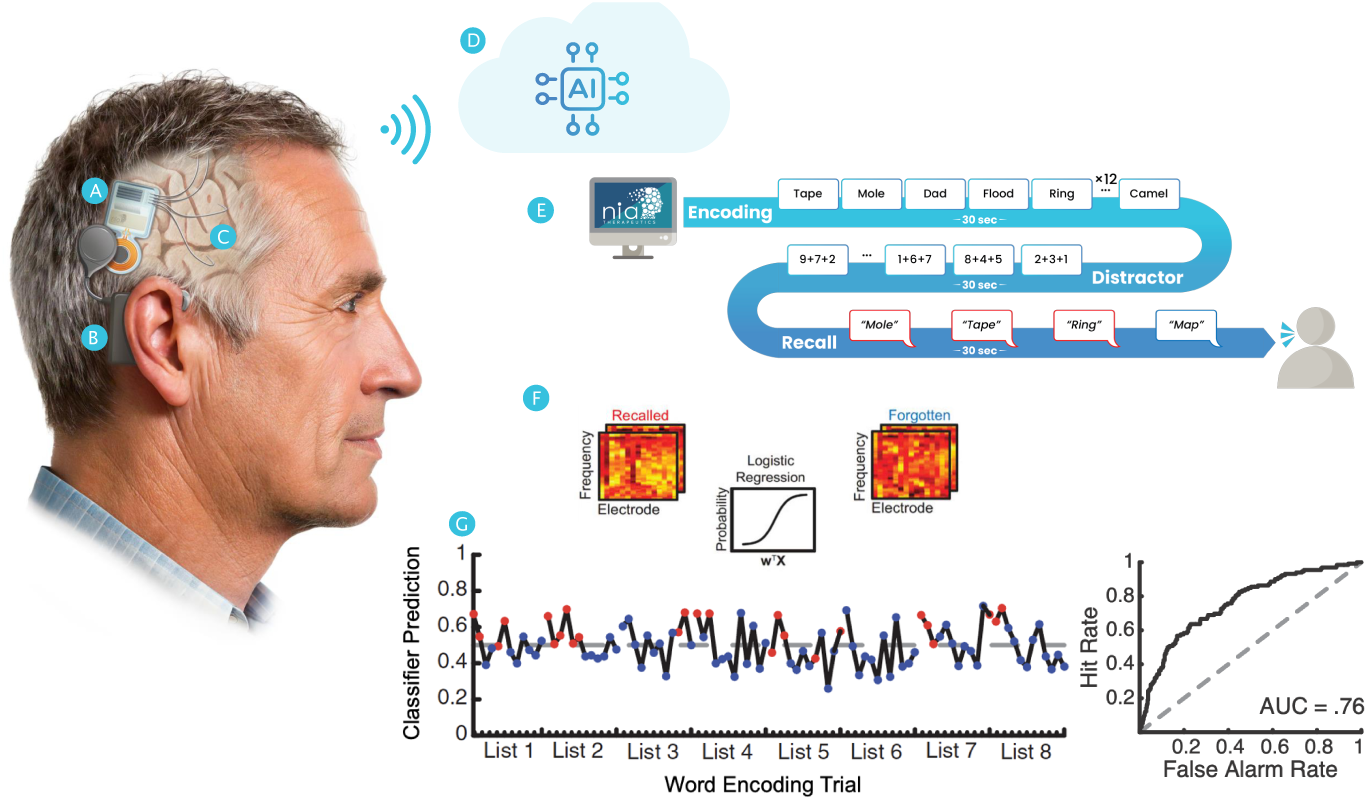
The Smart Neurostimulation System. Depiction of the SNS installed in an artist’s rendering of a typical patient. The SNS includes: (a) a cranial implant responsible for neural sensing and stimulation; (b) a wearable External Processor providing power and acting as a communications hub; (c) four depth leads with a total of 64 contacts; and (d) a cloud-based AI platform that personalize therapy for each patient. To predict momentary memory lapses, researchers record from widespread electrodes as subjects study and recall word lists (e). Multivariate classifiers (penalized logistic regression) learn a mapping between the distribution of spectral power across electrodes and the subsequent recall status of encoded items (f). These models can predict, in hold-out sessions, which list items subjects will remember (red dots) or forget (blue dots) (g). The SNS uses these models to trigger stimulation during predicted memory lapses and to optimize stimulation parameters.

During therapy personalization, the SNS records multi-channel field potentials while patients study and recall word lists (as in standard neuropsychological assessments, Figure 1E). Applying spectral filters to the EEG data enables the IPG to estimate LFP power at frequencies from 3-180 Hz. Machine-learning models trained on these spectral features predict mnemonic success (Figure 1F) and drive later closed-loop stimulation decisions during therapy (Figure 1G).

Figure 2 shows the IPG, EP, and Depth Leads. The IPG senses local field activity from 60 electrodes (plus four references), processes the data and controls therapy delivery. The EP inductively powers the IPG and relays data over a 2.2 Mbit/s link, sufficient to stream (500-Hz sampled) signals from every channel, with reliable transmission across 4–10 mm of scalp.

**Figure 2:**
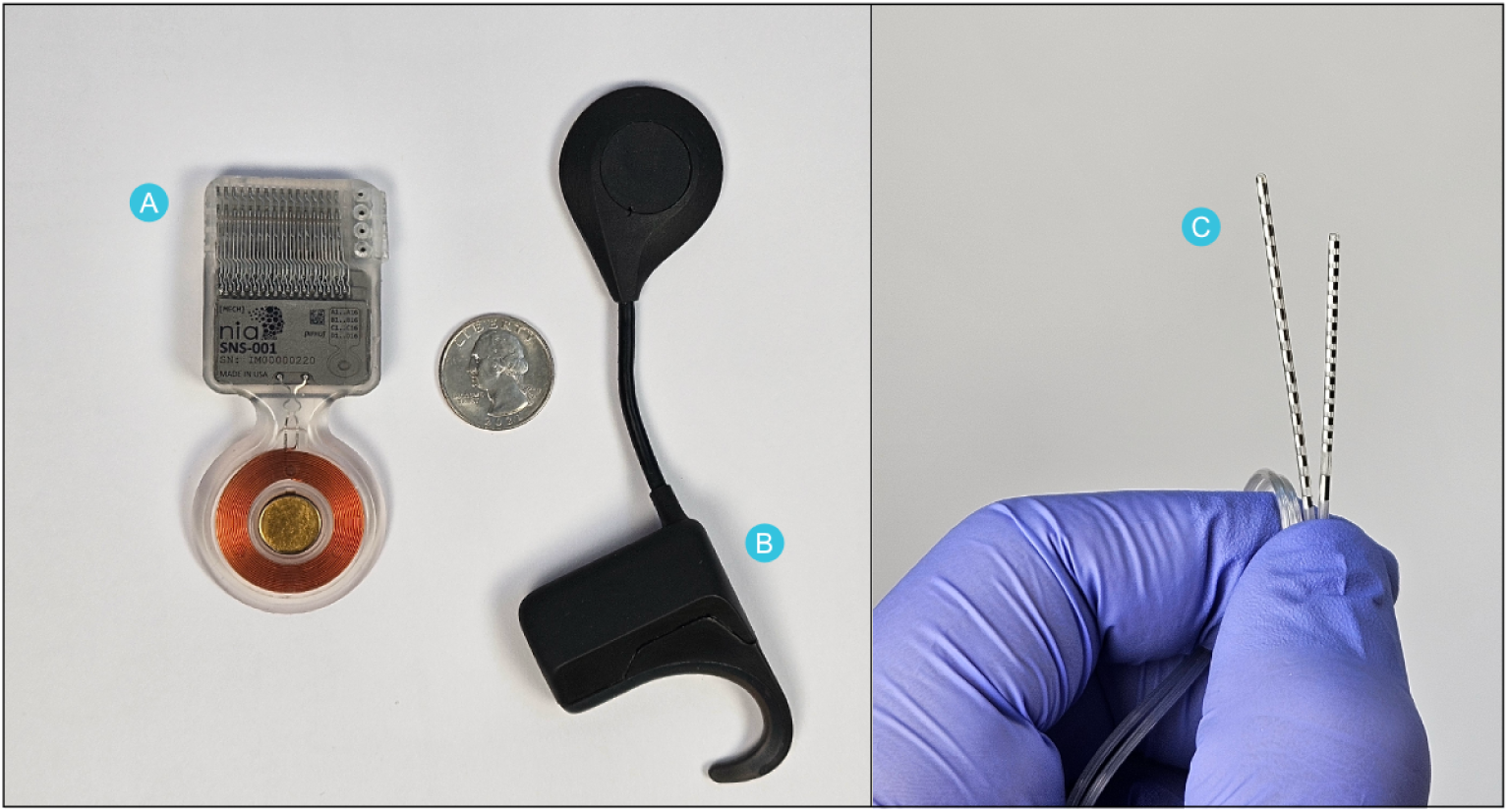
The Smart Neurostimulation System prototype, including a) the implantable pulse generator; b) four depth leads containing up to 16 electrodes each; and c) the external processor. All device materials were selected for biocompatibility and prior medical device usage. The silicone-encapsulated IPG houses a hermetically sealed titanium can surrounding the embedded electronics, platinum/iridium, stainless steel and Pellethane in the lead connector assembly, and Parylene-C surrounding the gold-coated magnet. The depth lead utilizes platinum/iridium electrodes, a stainless-steel set screw block, and a pellethane lead body. EP materials include ABS plastic, nylon and silicone. A U.S. quarter illustrates the scale of the device.

The EP serves as the system’s communication hub, providing USB for high-speed communication to the Programmer and Bluetooth low energy (BLE) for low-bandwidth mobile-app telemetry. The SNS end-to-end authenticates and encrypts all commands and responses using 128-bit AES-GCM, ensuring only authenticated users and devices can connect to the device, and preventing administration of malicious therapy parameters. As with cochlear implants, users recharge the EP overnight, outside therapy hours.

The SNS design arose from a need to meet three constraints: (i) recording field potentials from *>*32 electrodes in widespread brain locations at sampling rates *≥* 500 Hz (to discern high-frequency signals that predict mnemonic function [17, 18]); (ii) delivering rapid stimulation bursts triggered by onboard machine-learning models that predict memory lapses; and (iii) fitting within a cranial form factor proven safe in prior applications (e.g., the NeuroPace RNS).

Proof-of-concept studies [7, 19, 12] used bi-phasic square wave pulses at frequencies up to 200 Hz, amplitudes up to 3 mA, and durations up to 1 sec. These parameters, along with the requirement of inductive power transmission to avoid surgical battery replacement, require a capacity of 210mAh (this includes budgeting for inductive power loss and EP power usage).

The EP and IPG interact via an inductive link composed of two conductive coils (see Figure 3A,B). Command and response data flow bidirectionally through this link, while the IPG simultaneously transmits neural data to the EP. The EP can then send neural data to a PC over USB or to a mobile phone via BLE. A microcontroller (MCU) in the EP controls most of the system logic, including communication, safety systems, and firmware updates. A field programmable gate array (FPGA) controls the inductive link communication protocol on both the EP and the IPG. Power management hardware on the IPG regulates power input, while MCUs on the IPG and EP process all communication messages and manage therapy safety.

Figure 3C illustrates the electrical interface between IPG and neural tissue. The IPG lead interface holds four leads with 16 electrodes each. Sixty of the 64 platinum-iridium electrodes can sense and stimulate neural tissue; one electrode on each lead serves as an electrical reference. On the IPG, a proprietary NeuroModulation Integrated Circuit (NMIC) chip connects to the electrode contacts through DC-blocking capacitors. This custom Application-Specific Integrated Circuit (ASIC) was designed specifically for brain interfaces to provide sensing and stimulation capabilities [20, 21]. Inside the NMIC, there are 64 independent sensing and stimulation channels, one for each electrode contact. We model the brain-NMIC interface as an R-C circuit. Board-level filtering capacitors shunt noise to ground contacts. The IPG delivers charge-balanced biphasic pulses through any electrode pair on the same lead, sourcing current during the first phase and sinking it during the second.

Firmware architecture (Figure 4) supports command and control, battery management, firmware updates, event logging, and over-the-air updates. The firmware supports two modes: (i) therapy personalization and (ii) therapy delivery. During therapy personalization, the firmware transmits data from the IPG through the EP to an external PC (see *Streaming Path* Figure 4). In this mode, a clinician programmer initiates stimulation for adverse effects testing, therapy optimization, and therapy testing. During therapy delivery, the IPG senses neural data, classifies the brain’s state, and decides whether to apply stimulation. This is shown in Figure 4 as the *Therapy Delivery* path. In this path, FPGAs periodically process data, estimating spectral power within multiple frequency bands in a given time window (Figure 4A). A classifier in the MCU makes therapy decisions based on these features and the classifier-model parameters. If the classifier indicates that therapy should be applied, the IPG applies electrical stimulation to the brain. Configurable therapy parameters include target electrode, amplitude, frequency, pulse width, and duration.

**Figure 3:**
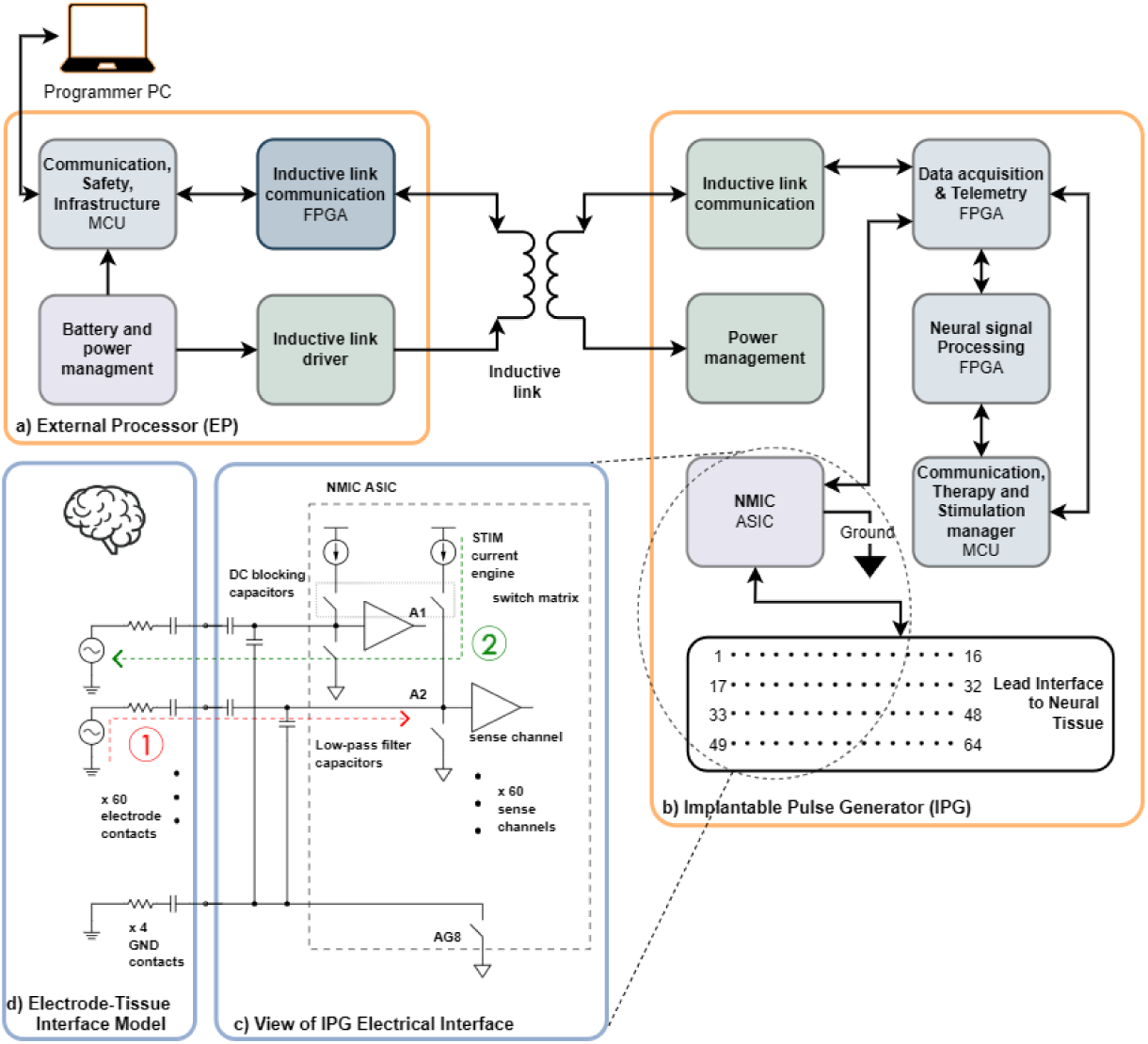
The SNS system hardware and electrical interface schematic. a) the External Processor (EP), and b) the Implantable Pulse Generator (IPG). c) Expanded view of the electrical interface between the stimulation/sensing ASIC (Neuromodulation Integrated Circuit; NMIC) and the leads. d) the Electrode-Tissue interface model. Information (data or commands) can be sent via the EP (a) to the IPG (b) and in reverse. The IPG records neural data via (c) and transmits it to the EP (which can then communicate with an external program (PC). The electrical interface and Electrode-Tissue interface (c) and (d) illustrates both sensing (1) and stimulation (2) circuit models.

**Figure 4:**
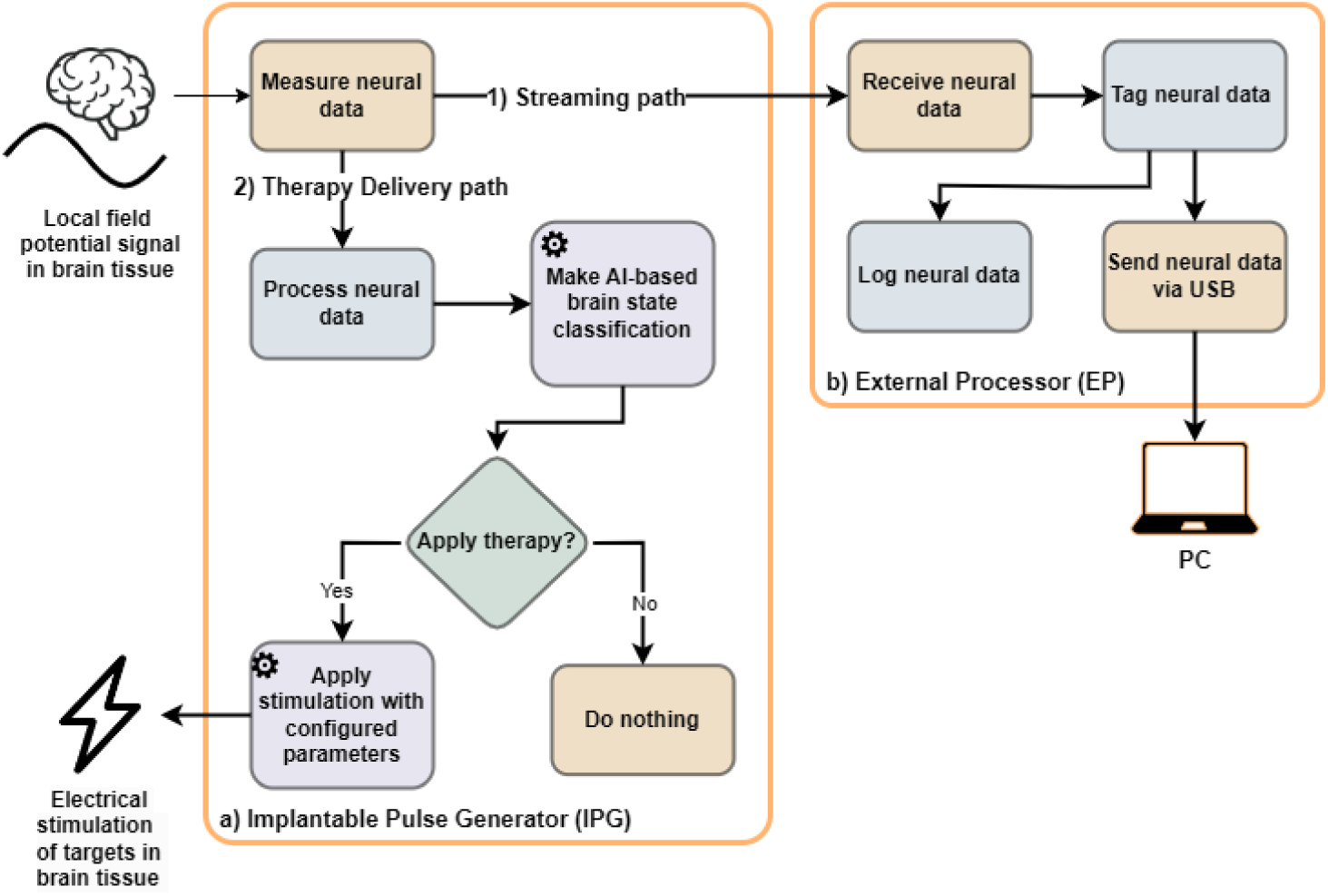
SNS firmware overview. The SNS includes firmware running on both the IPG (a) and the External Processor (b) to sense neural data and administer electrical stimulation. Once it is sensed from the brain, neural data can take two paths (numbered): 1.) a Streaming path where data is sent to an external computer via USB; and 2.) a Therapy Delivery path where the data are processed and therapy is administered based on brain state classification. Brain state classification and therapy delivery blocks are programmable and adaptable (marked with a gear icon).

## 3. Preclinical Ovine Study

To evaluate the performance of the Smart Neurostimulation System (SNS), we conducted a preclinical study using an ovine model. Figure 5 outlines the study design. After receiving approval from the Institutional Animal Care and Use Committee (IACUC), we performed CT imaging to guide trajectory planning for electrode implantation. On Day 0, a functional neurosurgeon (R.E.G. or B.C.L.) implanted a single SNS IPG and two depth leads per animal, followed by post-operative CT scans to confirm lead placement. Each animal was allowed a six-day recovery period before initiating neural recordings and stimulation.

Animals were housed under veterinary supervision and allowed to move freely within their enclosures throughout the study. At the study’s conclusion, each animal was humanely euthanized. Table 1 summarizes the characteristics of each animal and the implanted devices.

**Figure 5:**
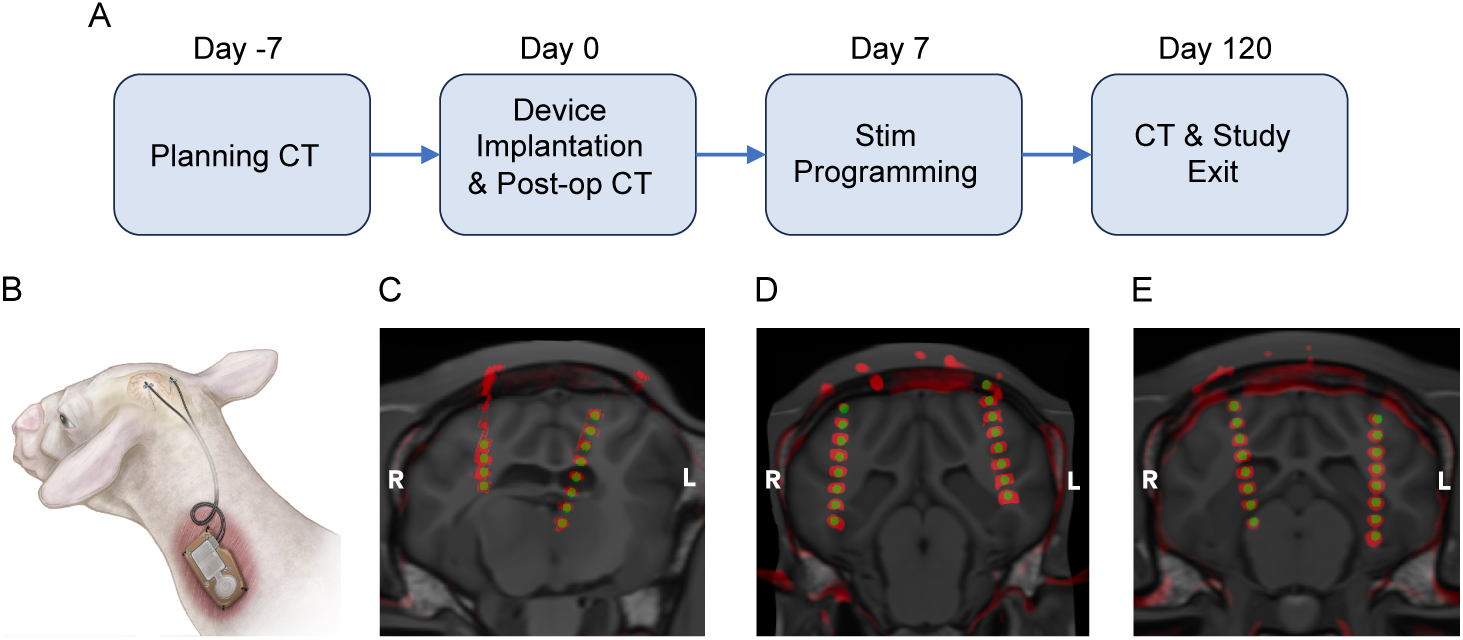
Preclinical study design and surgical approach. (A) Time course of the preclinical study. CT imaging was collected on Day -7, Day 0 (post-implantation) and Day 120 (prior to study exit). (B) The IPG and a protective exoskeleton were implanted into a pocket in the neck, and two leads were implanted into the left and right parietal lobes. An EP device bandaged on the skin overlying the IPG coil (not shown) communicated with and powered the IPG. (C, D, E) CT imaging showing the locations of the depth leads in animals S001, S002 and S003, respectively.

To localize the electrodes anatomically, we registered post-operative CT scans to the Turone sheep brain MRI atlas [22, 23] using ITK-SNAP. The iterative metal artifact reduction (iMAR) algorithm was applied to reduce CT signal distortion from the electrodes. We used bright metal artifacts to identify electrode positions and extracted 3D coordinates relative to anatomical regions defined by the atlas. Table 2 reports the anatomical locations for all monopolar electrodes.

Each SNS lead included eight channels, numbered with even integers from 2 to 16. Channel 8 served as the sensing reference, and all reference electrodes were internally connected within the IPG. (The commercial control lead did not include a reference electrode and thus did not contribute to the reference scheme in animal 1.) For analysis, we generated bipolar “virtual” recordings by subtracting voltages between adjacent non-reference electrodes (e.g., A2–A4). Bipolar referencing, computed as the voltage difference between pairs of adjacent electrodes, filters out signals common to both channels, thereby improving signal-to-noise ratio. This referencing scheme attenuates oculomotor and electromyographic artifacts that can mix with neural signals, and large-*N* human studies have shown that bipolar referencing of in-traparenchymal electrodes better resolves the neural correlates of memory than average referencing [24]. On each SNS lead, the fourth most distal electrode (denoted number eight) served as the electrical reference for the other electrodes.

**Table 1:**
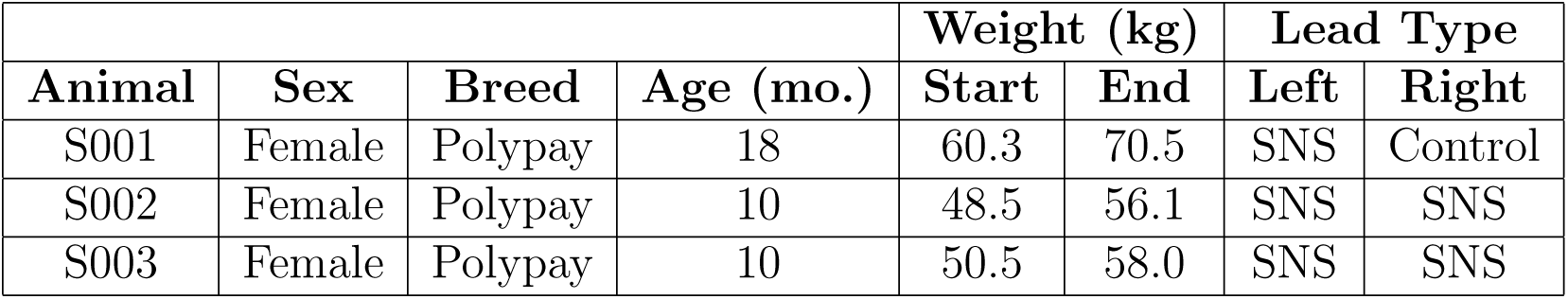
Animal characteristics and depth leads implanted into left and right hemispheres. All animals had one SNS IPG implanted at the bottom of the neck and one depth lead placed into each hemisphere. In animal S001, one 8-channel SNS depth lead was placed in left hemisphere and a Medtronic Control lead (model 3387S-40) was placed in right hemisphere. All other animals had SNS 8-channel depth leads placed into each hemisphere.

|  |  |  |  | Weight (kg) |  | Lead Type |  |
| --- | --- | --- | --- | --- | --- | --- | --- |
| Animal | Sex | Breed | Age (mo.) | Start | End | Left | Right |
| S001 | Female | Polypay | 18 | 60.3 | 70.5 | SNS | Control |
| S002 | Female | Polypay | 10 | 48.5 | 56.1 | SNS | SNS |
| S003 | Female | Polypay | 10 | 50.5 | 58.0 | SNS | SNS |

All three animals underwent surgical implantation without intraoperative complications. The most observed adverse effect was localized swelling at the IPG site, which resolved with the application of a pressure bandage. Routine veterinary assessments confirmed normal wound healing, and there was no evidence of infection, hemorrhage, or tissue damage at the lead implantation sites. One serious adverse event occurred in Animal 3: erosion of the skin overlying the IPG coil. This was attributed to the EP device used in this animal exerting higher-than-intended pressure on the skin, leading to the early euthanasia of the animal (day 61).

## 4. Using Neural Recordings to Predict Movement

We first asked whether the SNS could record and decode neural signals related to animal behavior. With a triaxial accelerometer sampling at 10 Hz, we gauged the animal’s activity level throughout two-hour sessions. We selected non-overlapping 1-second epochs from longer sustained periods of movement or stillness, and used the neural data from these epochs to classify movement in hold-out sessions. Whenever an accelerometer reading exceeded 0.03*m/s*^2^, if *>* 30% of samples in the following 10 seconds exceeded the accelerometer threshold, we created a “movement” epoch. Whenever an accelerometer reading was below 0.03*m/s*^2^, if all samples in the prior 60 and subsequent 15 seconds were below the threshold, we created a “stillness” epoch. We included sessions that had at least 100 movement and 100 stillness epochs, and neither class made up more than 80% of the total epochs.

**Table 2:** Monopolar electrode locations. Labels A and B refer to Nia eight-channel leads implanted in the left and right hemispheres, respectively; Label M refers to the Medtronic control lead implanted in S001. Localization methods appear in the text.

| Subject | Electrode Label | X | Y | Z | Hemisphere | Anatomical Label |
| --- | --- | --- | --- | --- | --- | --- |
| S001 | A2 | -0.5 | 5.5 | 2 | L | Nucleus intermediodorsalis |
|  | A4 | -1.75 | 5.5 | 6 | L | Nucleus habenulae |
|  | A6 | -2.75 | 5.5 | 8.75 | L | Fornix |
|  | A8 | -4 | 6 | 12 | L | Lateral ventricle |
|  | A10 | -5 | 6 | 15.25 | L | White matter bundle |
|  | A12 | -6 | 6 | 18.25 | L | Lateral gyrus |
|  | A14 | -6.75 | 6.25 | 21.5 | L | White matter bundle |
|  | A16 | -8 | 6.25 | 25 | L | White matter bundle |
|  | M1 | 13.25 | -1.25 | 10 | R | White matter bundle |
|  | M2 | 13.75 | -1.25 | 12.5 | R | Caudate |
|  | M3 | 14.25 | -1.25 | 15.25 | R | White matter bundle |
| S002 | A2 | -19.25 | -2.5 | -31.75 | L | White matter bundle |
|  | A4 | -18.5 | -2.5 | -28.25 | L | White matter bundle |
|  | A6 | -18 | -2.5 | -24.75 | L | Suprasylvius gyrus |
|  | A8 | -17.25 | -2.5 | -21.25 | L | Suprasylvius gyrus |
|  | A10 | -16.75 | -2.5 | -18 | L | White matter bundle |
|  | A12 | -16.25 | -2.5 | -14.75 | L | Suprasylvius gyrus |
|  | A14 | -15.5 | -2.5 | -11 | L | Suprasylvius gyrus |
|  | A16 | -14.75 | -2.5 | -7.75 | L | Skull |
|  | B2 | 18.5 | -2.5 | -37.25 | R | Temporal gyrus |
|  | B4 | 18.25 | -2.5 | -33.75 | R | White matter bundle |
|  | B6 | 17.75 | -2.5 | -30 | R | white matter bundle |
|  | B8 | 17.5 | -2.5 | -26.5 | R | Posterior sylvian gyrus |
|  | B10 | 17 | -2.5 | -23 | R | White matter bundle |
|  | B12 | 17.25 | -2.5 | -19.25 | R | White matter bundle |
|  | B14 | 16.75 | -2.5 | -15.75 | R | White matter bundle |
|  | B16 | 16.25 | -2.5 | -12.75 | R | Suprasylvius gyrus |
| S003 | A2 | -16 | -3.25 | -3 | L | Parahippocampal cortex |
|  | A4 | -16 | -3.25 | 0.25 | L | Hippocampus |
|  | A6 | -16 | -3.25 | 3.75 | L | Hippocampus |
|  | A8 | -16.25 | -3.25 | 7.5 | L | White matter bundle |
|  | A10 | -16.25 | -3.25 | 10.75 | L | White matter bundle |
|  | A12 | -16.5 | -3.25 | 14.25 | L | White matter bundle |
|  | A14 | -16.25 | -3.25 | 17.75 | L | White matter bundle |
|  | A16 | -16 | -3.25 | 21.25 | L | Suprasylvius gyrus |
|  | B2 | 9.5 | -3.25 | 0 | R | White matter bundle |
|  | B4 | 10 | -3.25 | 3.25 | R | White matter bundle |
|  | B6 | 10.5 | -3.25 | 6.75 | R | Hippocampus |
|  | B8 | 11.25 | -3.25 | 10.25 | R | Hippocampus |
|  | B10 | 11.75 | -3.25 | 13.75 | R | White matter bundle |
|  | B12 | 12.25 | -3.25 | 17.25 | R | White matter bundle |
|  | B14 | 13 | -3.25 | 20.5 | R | White matter bundle |
|  | B16 | 13.5 | -3.25 | 24 | R | Ectolateral gyrus |

The final dataset comprised 48, 36, and 48 sessions from the three sheep, respectively. For each one-second epoch, we calculated the spectral powers using Morlet wavelets at eight log-spaced frequencies, ranging from 6 to 180 Hz. The powers were then log-transformed and z-scored within each recording channel and frequency, and then averaged across the one-second epoch. The spectral powers for every epoch at each frequency and channel served as the features input to an *L*^2^-penalized logistic regression classifier, where the labels indicated the movement or stillness identity of each epoch. Using an 80-20 session split, we trained the classifiers to discriminate brain activity predictive of movement and stillness. For cross-validation, we randomly repeated the 80-20 session split, with the number of permutations equal to the number of sessions. We computed the area under the receiver operating characteristic curve (AUC) to quantify classifier performance for each permutation.

Figure 6A demonstrates the cross-validation in an example session. Accelerometer readings, in cyan, suggest the animal exhibited two periods of reliable stillness and two periods of significant activity. Red and blue dots indicate classifier outputs for epochs labeled as “movement” and “stillness”, respectively. Blue dots tend to fall under the 0.5 classifier output threshold, and red dots tend to stay above it, indicating that classifier predictions agreed with the accelerometer reading interpretations in most cases.

We illustrate receiver operating characteristic (ROC) curves in Figure 6B. For each animal, the hit rate rises quickly towards 1.0 as the false alarm rate increases, suggesting neural features reliably classify the animal’s activity in holdout data. Across the cross-validation test partitions, we found AUC values of (*M* = 0.924*, SD* = 0.031), (*M* = 0.979*, SD* = 0.008), and (*M* = 0.969*, SD* = 0.008) for the three sheep.

Previous analyses of neural data during motor tasks have identified increases in high-frequency power (*>* 30 Hz) and decreases in low-frequency power [17] during and immediately preceding movement. To determine whether our classifier uncovered similar neural correlates of movement, we applied the Haufe method [25] to the weights obtained from the optimal classifier fit to cross-validation training datasets. This method transforms classifier weights to account for the covariances between features. In Figure 6C, we illustrate that increased high-frequency power and decreased low-frequency power tended to mark high activity periods. Although aggregating across all electrodes illustrated a similar trend across animals, the classifiers identified the unique pattern of neural activity related to behavior within each animal.

**Figure 6:**
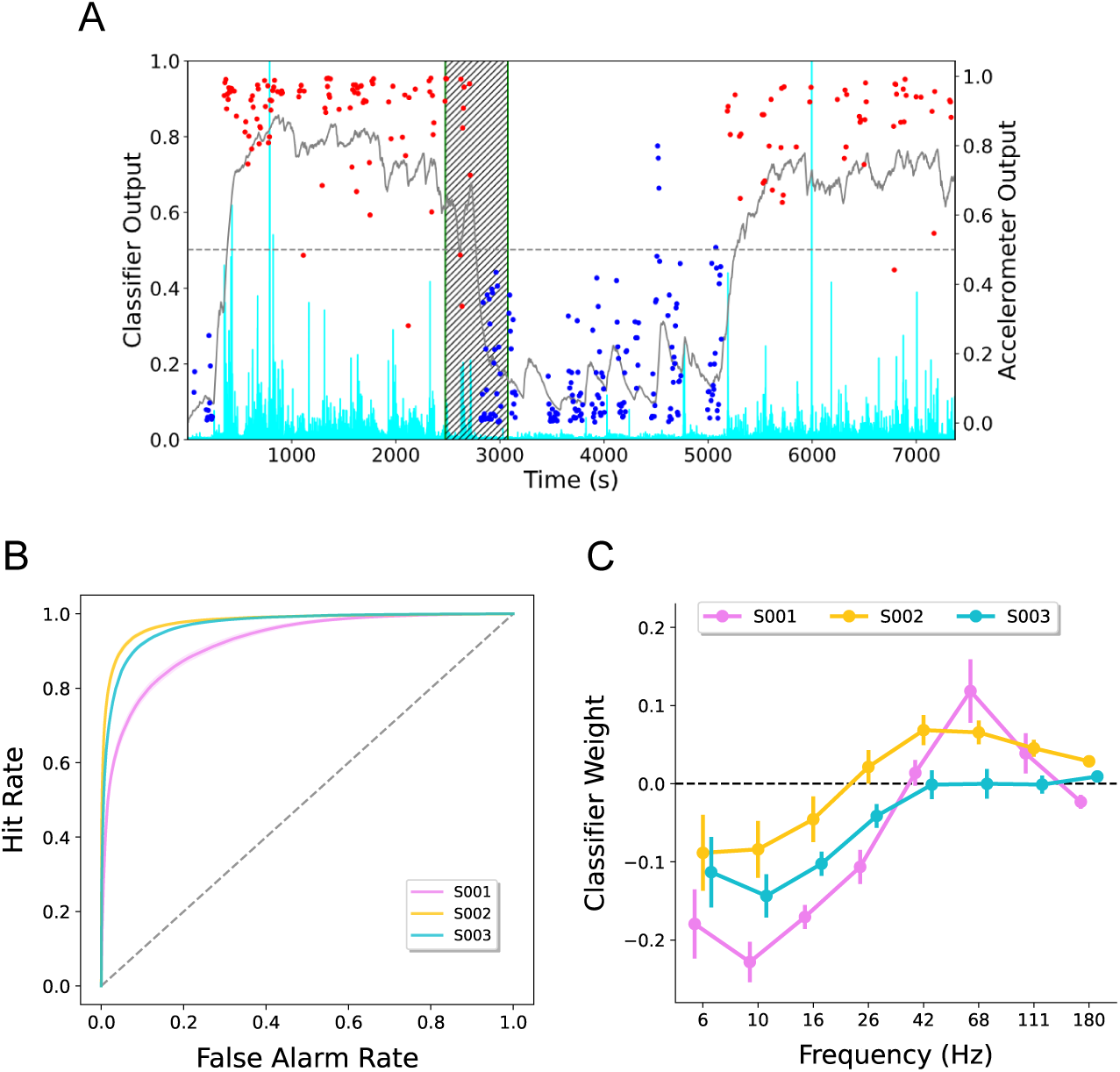
Classification of animal movement. (A) We trained an L2 logistic regression classifier to predict periods of movement vs. stillness based on spectral features. Across a two-hour hold-out session, we show that classifier output probability (gray) tracks periods of movement (red) and stillness (blue). These representative periods were chosen based on the accelerometer magnitude readings (cyan) sampled at 10Hz and high-pass filtered with a cutoff at 0.1Hz. (B) A ROC analysis, which shows the relation between true and false positives as a function of decision criterion, quantifies classifier performance. Error bars represent one standard error of the mean across cross-validation permutations. (C) The Haufe method reveals the degree of influence of different features on classification performance while adjusting for the covariance structure in the model. Here we see that increased weights on high frequencies and decreased weights on low frequencies predicted movement. Error bars represent one standard error of the mean across recording channels.

## 5. Neural Recordings and Stimulation Performance

To evaluate the SNS’s ability to modulate neural activity, we ran two stimulation experiments. We compared spectral power in the 200–950 ms post-stimulation period to power in the -750–0 ms pre-stimulation period.

Analyzing the post-stimulation period, isolated stimulation induced brain activity that persists beyond the stimulation interval, and the 200-ms post-stimulation buffer attenuated potential stimulation artifact. Guided by prior work on spectral biomarkers of cognition, we focused on alpha (8–12 Hz) and gamma (78–82 Hz) band power. We used Welch’s method to calculate the power spectral densities during the pre- and post-stimulation periods, which does not require a buffer that risks leakage from the stimulation period. Before further analysis, we log-transformed each power value, as in previous work [26, 17], and calculated the z-scores of spectral powers within each frequency, channel, and session. For every stimulation event, *i*, within each frequency, *f*, and channel, *c*, we computed the difference in post- and pre-stimulation power as Δ_P(f,c,i)_ = P(f, c, i)_*post*_ − P(f, c, i)_*pre*_ (Figure 7C).

### 5.1. Parameter Search: Experiment 1

Experiment 1 tested whether the SNS could reliably modulate neural activity. Applying stimulation at varying amplitudes and frequencies at a single pair of neighboring electrodes, we evaluated power changes at the remaining non-anode, non-cathode electrodes on the same implanted lead. On each trial, the SNS applied 300 *µ*s biphasic stimulation pulses at frequencies of 25, 50, 100 and 200 Hz and amplitudes of 0 (sham), 0.2, 0.4, 0.6, 0.8, and 1 mA across a bipolar pair of neighboring electrodes (*A*10 *− A*12) in left frontal white matter (Figure 5D). To minimize order effects, we shuffled stimulation *trials* to create 10 uniquely ordered session protocols, each with 30 trials of each of the 24 frequency *×* amplitude combinations. Each trial applied stimulation for 500 ms, followed by a 2.0–2.25 s inter-trial interval (Figure 7A). One sheep completed 124 sessions.

Increasing stimulation amplitude and frequency boosted both alpha and gamma power (Figure 8). These effects broadly align with prior stimulation parameter search experiments in humans [19]. We evaluated these trends with a linear mixed effects model containing fixed effects for stimulation amplitude, frequency, and their interaction, a random intercept for session, and had predictor variables centered and normalized to the range -1 to 1. Our model identified statistically significant linear effects of both stimulation amplitude *β* = 0.311*, CI* = [0.303, 0.319] and frequency, *β* = 0.140*, CI* = [0.134, 0.147] on stimulation-related changes in alpha power. Our model also identified a significant interaction between amplitude and frequency, *β* = 0.159*, CI* = [0.149, 0.168], as seen in the increasing strength of the amplitude-power relation for increasing stimulation frequency evident in Figure 8. We observe significant relations of stimulation amplitude, *β* = 0.013*, CI* = [0.005, 0.022], frequency, *β* = 0.032*, CI* = [0.025, 0.040], and their interaction, *β* = *−*0.011*, CI* = [*−*0.022*, −*0.001], with changes in gamma power. These results align with previous work finding post-stimulation increases in high-frequency activity (HFA) [19]. Prior research has shown that HFA increases correlate with increases in neural firing rate, linking stimulation-related spectral changes to the modulation of neural activity [27].

**Figure 7:**
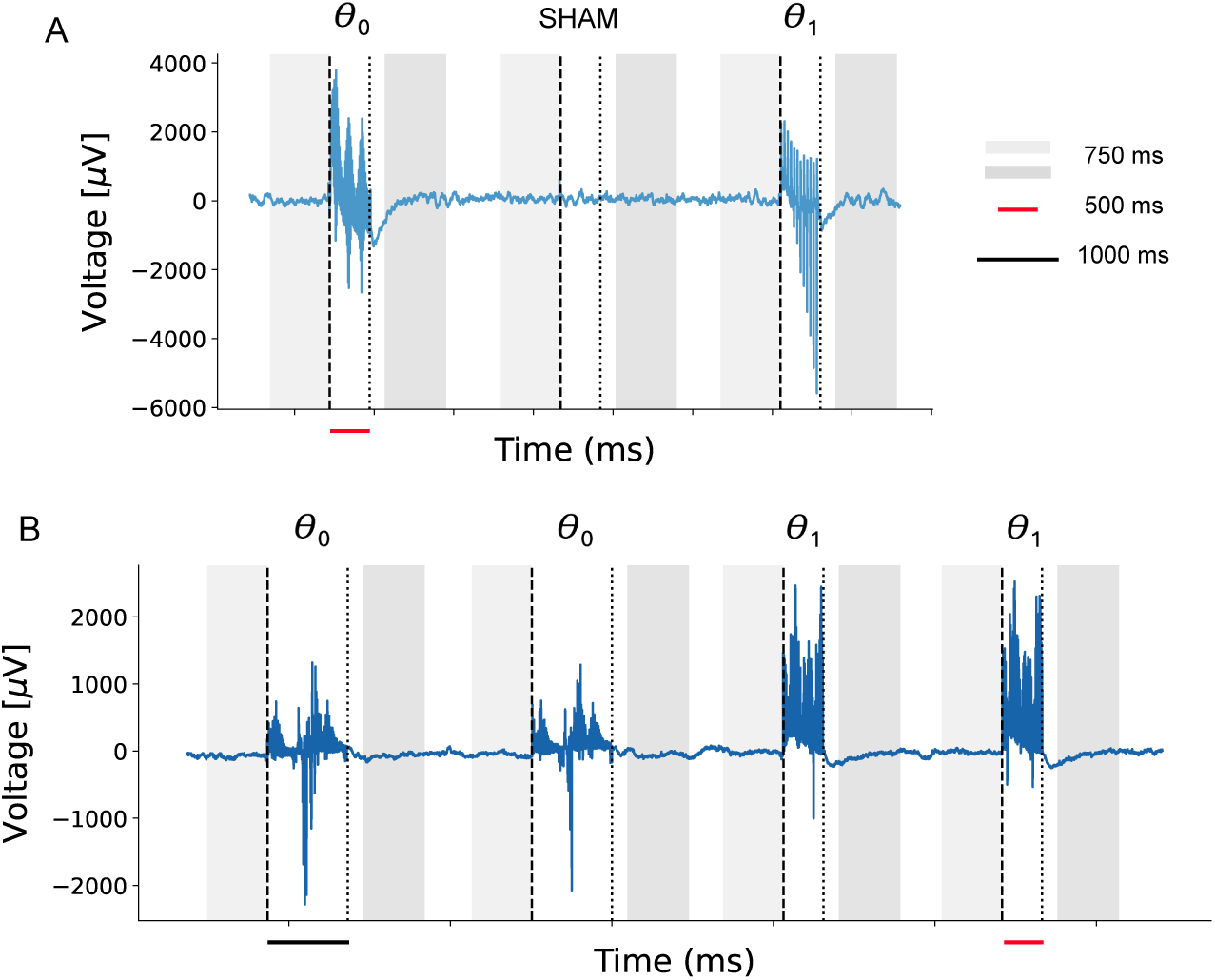
Parameter search methods. We applied stimulation with varying parameter sets *θ* and analyzed spectral powers from -750 to 0 ms before stimulation onset and 200 to 950 ms following stimulation offset. (A) In Experiment 1, the SNS delivered 500 ms bi-phasic stimulation bursts at varying amplitudes [Sham, 0.2, 0.4, 0.6, 0.8, 1 mA] and frequencies [*f* = 25, 50, 100 and 200 Hz] at a single target location (A10-A12), with the parameter set updating each stimulation event. Example field potential recorded from a bipolar electrode pair (A14-A16). (B) In Experiment 2, the SNS delivered bi-phasic stimulation bursts at varying amplitudes [0.5, 1, 1.5 mA], frequencies [100, 200 Hz], and durations [500, 1000 ms] at twelve target locations, with the parameter set updating every two stimulation events. Example field potential recorded from a bipolar electrode pair (B4-B6).

**Figure 8:**
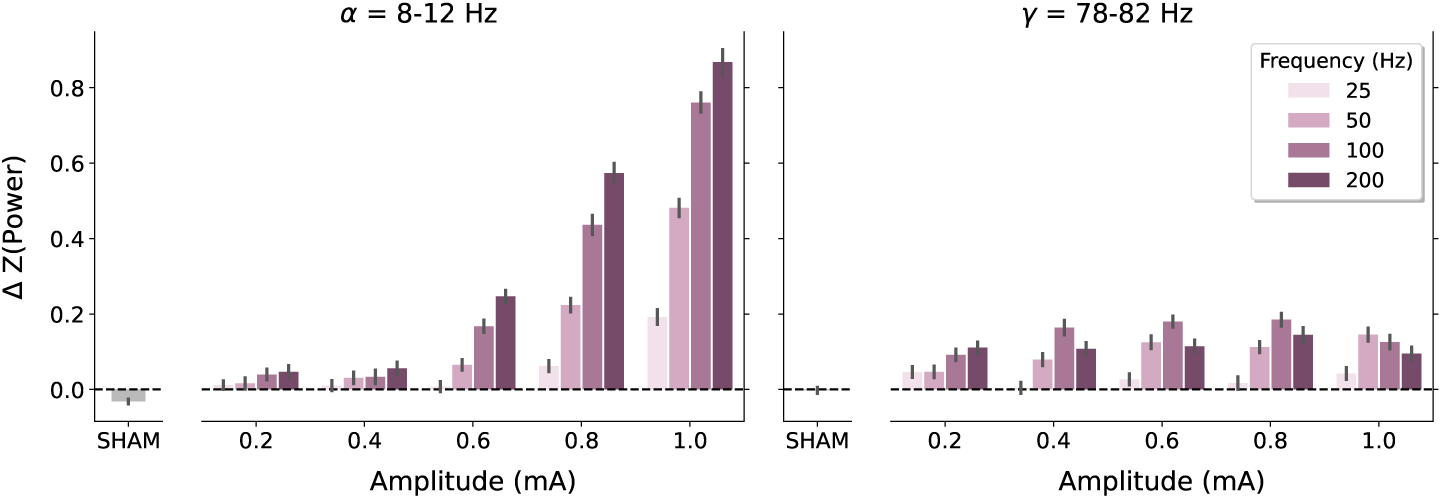
Stimulation’s physiological effects: Experiment 1. The effect of stimulation amplitude and frequency on alpha-band power (8-12 Hz, left panel) and gamma-band power (78-82 Hz, right panel). This analysis averages over the three electrode pairs that do not overlap with either the anode or cathode. Increasing stimulation amplitude and frequency results in greater power increases.

### 5.2. Parameter Search: Experiment 2

Experiment 2 further explored the modulatory capabilities of the SNS. In two animals, we stimulated 12 neighboring electrode pairs with varying amplitude, frequency, and duration. We evaluated stimulation’s effects on power at the remaining electrodes on the same implanted lead not involved in stimulation and the electrodes on the other implanted lead. On each trial, the SNS applied 300 *µ*s biphasic stimulation pulses at frequencies of 100 and 200 Hz, amplitudes of 0.5, 1, and 1.5 mA, and durations of 500 and 1000 ms. As illustrated in Figure 7B, stimulation trials occurred in blocks of two stimulation events with the same stimulation parameters. Within a block, the two stimulation events occurred with a random inter-trial interval of either 1000 *−* 1250 ms or 2000 *−* 2250 ms. Across blocks, there was a random 2000 *−* 2250 ms inter-stimulus interval. We collected a total of 42 and 154 sessions from the two sheep, respectively.

Figures 9-10 reveal considerable site-to-site heterogeneity in the effects of stimulation amplitude, frequency, and duration on alpha and gamma. Overall, stimulation-related power increases appeared far more prevalent than power decreases. To evaluate these effects, we fit linear mixed-effects models for each stimulation location in each animal (fixed effects: amplitude, frequency, duration, and pairwise interactions; random intercept for session; with predictors centered and normalized in Experiment 1). We FDR corrected across the parameters estimated within each subject and frequency band [28]. For some stimulation locations (e.g., B10-B12 in both animals), increasing stimulation amplitude and frequency led to greater increases in both alpha and gamma power (all *t >* 8 and all *p <* 0.001), mirroring Experiment 1. However, at other sites, we observe no effect of amplitude (e.g., A2-A4 in animal S002) or frequency (e.g., A4-A6 in animal S003) on alpha or gamma power. In some cases, we observe negative effects of amplitude (e.g., B4-B6 in animal S003, alpha and gamma power, both *t < −*4.0 and *p <* 0.001) but we observe no negative effects of frequency.

Figure 11 summarizes the modeling results. More than half of all stimulation targets increased alpha and gamma power, and only two consistently decreased alpha power. Increasing stimulation amplitude tended to consistently increase post-stimulation changes in alpha and gamma; however, we did observe a small number of locations where changes in power decreased with increasing amplitude. Stimulation frequency exhibited a similar pattern to stimulation amplitude. We detected many fewer stimulation locations where duration reliably modulated either alpha or gamma power, and those locations did not show a clear directional pattern. The pairwise interactions between amplitude, frequency, and duration exhibited a diversity of results, mirroring that seen in Figures 9 and 10.

## 6. Discussion

Brain-responsive stimulation has emerged as an established therapy for epilepsy and movement disorders, with promise for expanded indications including chronic pain and depression [2, 29, 3, 4, 5, 6]. However, FDA-cleared platforms only decode a single spectral feature from a handful of channels. The Smart Neurostimulation System (SNS) overcomes this limitation by measuring electrical fields across 60 bipolar contacts on four depth leads and providing embedded algorithms for spectral processing and classifier-based stimulation control.

The ability to stimulate the brain in response to distributed patterns of neural activity may be particularly useful for neurocognitive and affective disorders where functional impairments vary dramatically from moment to moment [30]. As proof-of-concept, Ezzyat and colleagues [8, 11, 12] deployed a closed-loop algorithm on a partially externalized device, showing that stimulating the lateral temporal cortex during predicted memory lapses produced significant memory improvements in sham-controlled, double-blinded studies of neurosurgical epilepsy patients. By training logistic regression classifiers on spectral activity during learning, these studies could reliably predict which items would be recalled or forgotten, then trigger stimulation during predicted memory lapses. Mnemonic benefits only appeared for closed-loop stimulation — random stimulation did not improve memory.

**Figure 9:**
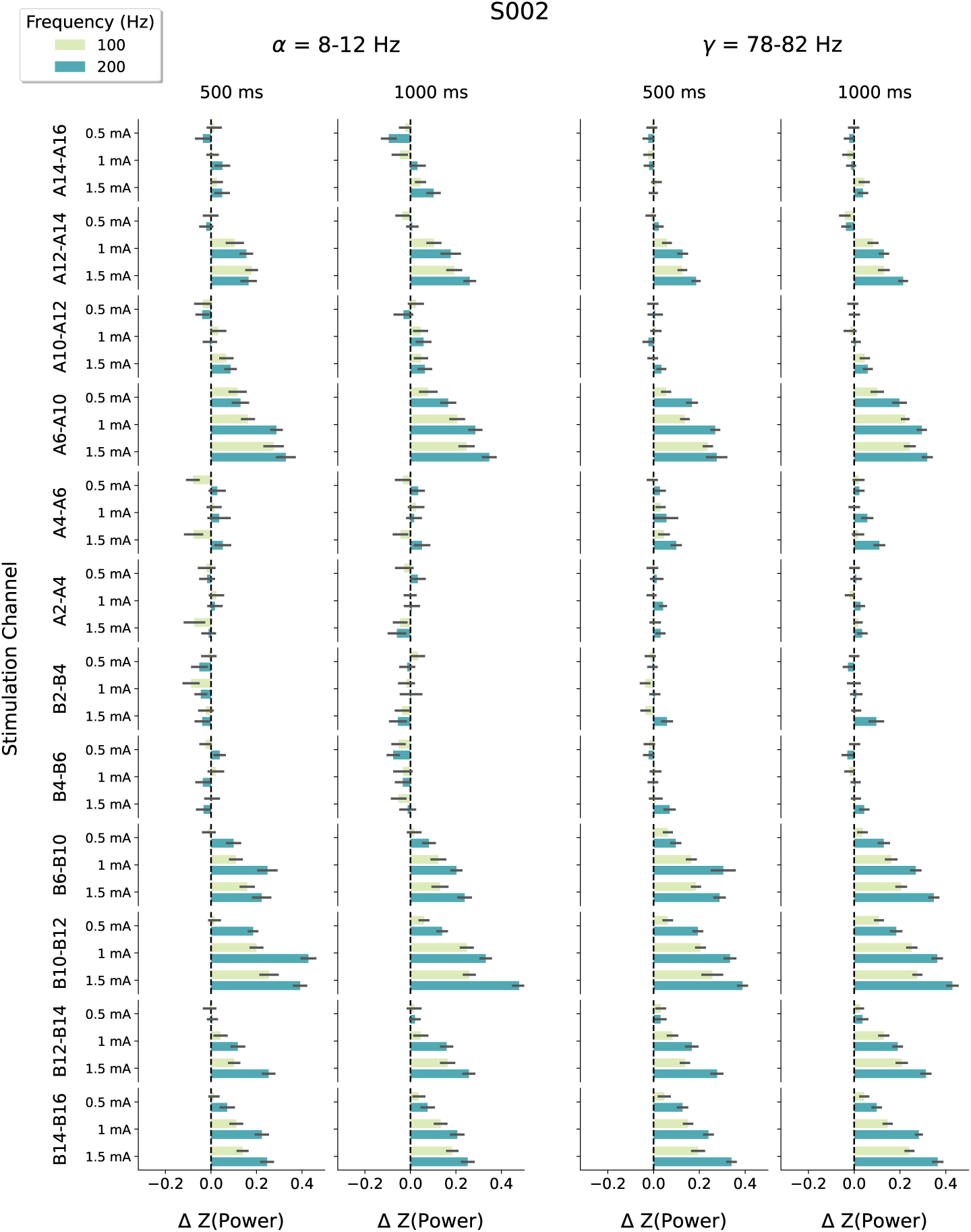
Stimulation’s physiological effects: Experiment 2. The effect of stimulation amplitude, frequency, duration, and location on alpha-band power (8-12 Hz, left panel) and gamma-band power (78-82 Hz, right panel) for animal S002. This analysis averages over the nine electrode pairs that do not overlap with either the anode or cathode. The effects of the stimulation parameters depend greatly on the location of stimulation.

**Figure 10:**
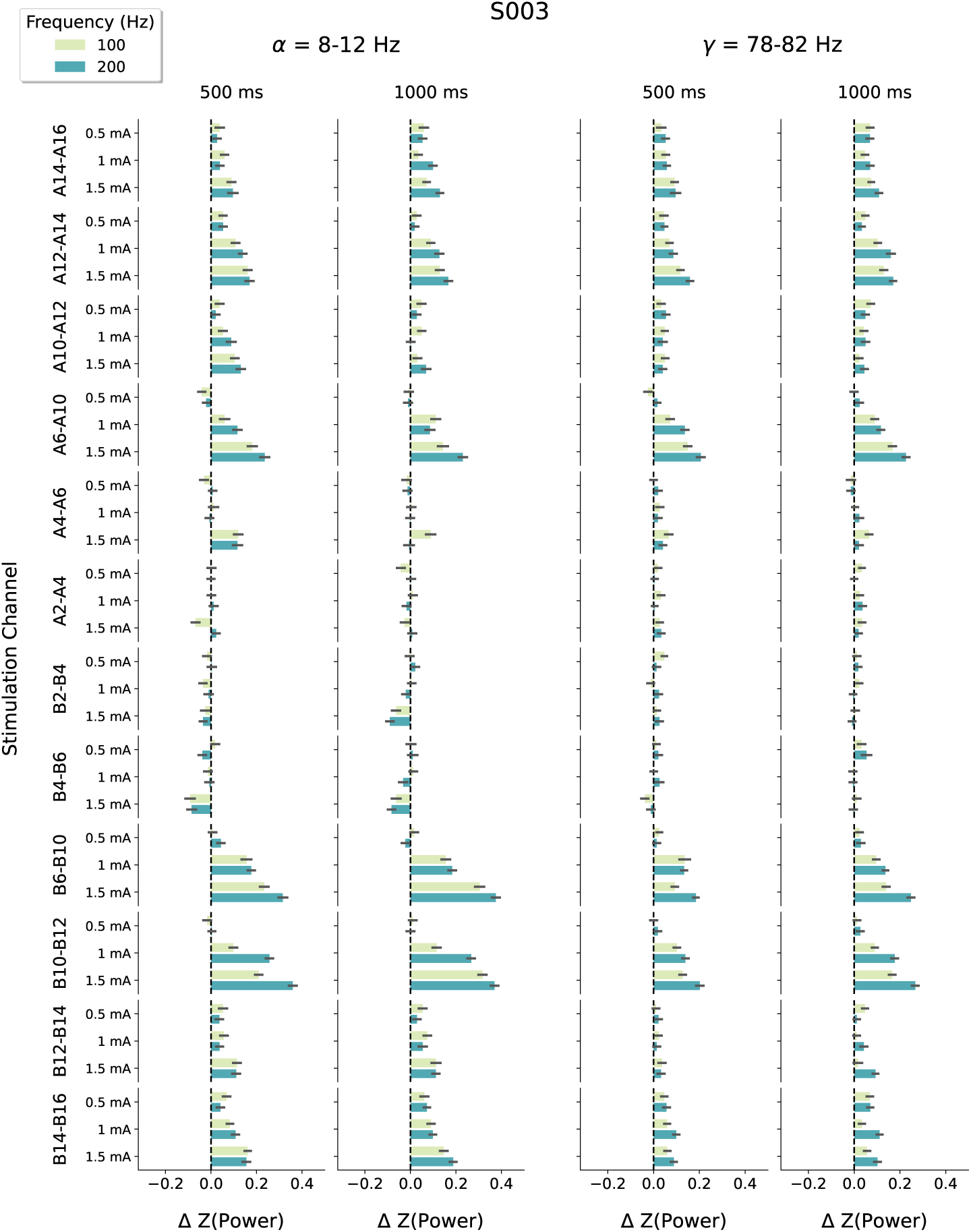
Stimulation’s physiological effects: Experiment 2. The effect of stimulation amplitude, frequency, duration, and location on alpha-band power (8-12 Hz, left panel) and gamma-band power (78-82 Hz, right panel) for animal S003. This analysis averages over the nine electrode pairs that do not overlap with either the anode or cathode. The effects of the stimulation parameters depend greatly on the location of stimulation.

**Figure 11:**
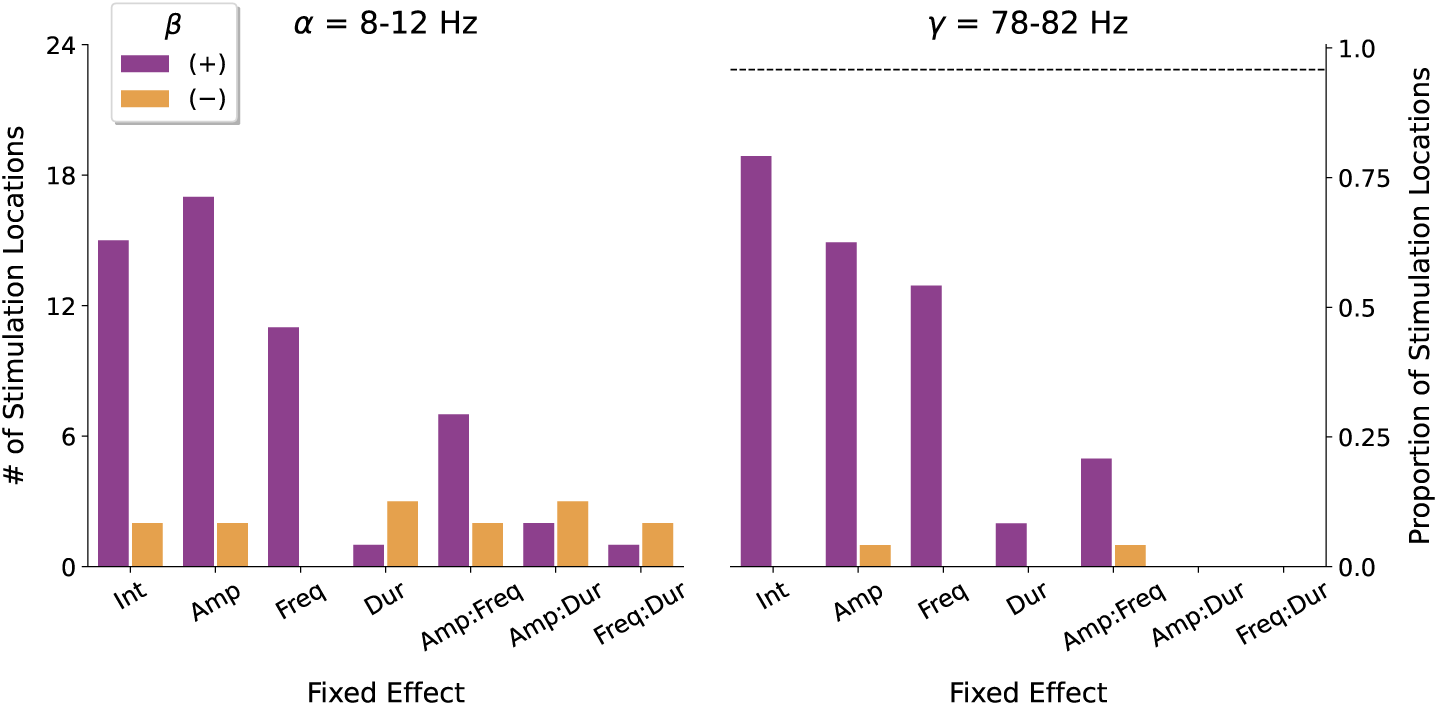
Linear mixed effects modeling of stimulation parameters. Number of stimulation targets exhibiting significant effects of stimulation amplitude, frequency, duration (and all pairwise interactions) on changes in alpha- and gamma-band power. Aggregated data from animals S002 and S003.

We designed the SNS to deliver closed-loop stimulation chronically via an AI-enabled brain implant (IPG) that communicates wirelessly with an external processor (EP). The EP, modeled on a cochlear implant sound processor, delivers power inductively to the IPG and acts as a communication hub with the cloud-based AI platform. Clinical programmer software enables patient-specific models to trigger therapeutic stimulation based on real-time brain state variability. In memory applications, predicted memory lapses will trigger stimulation, and post-minus-prestimulation classifier output will guide stimulation parameter optimization.

### 6.1. Comparison with Existing Systems

The NeuroPace RNS represents the closest predicate to the SNS. Both systems record field potentials and stimulate the brain in closed loop with onboard signal processing. However, the RNS records from a maximum of six channels in two brain regions versus the SNS’s 60 channels across four regions. Because seizures often localize to small brain regions, six channels significantly reduce seizures in many patients. However, memory-related brain signals occur across distributed networks [9, 10], requiring the widespread surveillance capabilities of the SNS and the ability to adaptively select among a larger set of stimulation targets. Analyses of large open-data sets [31, 32] informed the SNS design by identifying whole-brain patterns of neural activity that signal periods of good and poor memory.

The FDA cleared the Medtronic *Percept* adaptive DBS system for Parkinson’s disease in 2025. Percept records field potentials from each segmented STN or GPi lead, extracts a single *β*-band (13–30 Hz) power estimate once per second, and automatically raises or lowers stimulation amplitude when that biomarker crosses clinician-defined thresholds [2, 33]. In the pivotal trial, the algorithm maintained motor benefit while significantly reducing delivered charge. Despite these gains, Percept senses at most two bipolar channels and reacts to one spectral feature, making it unsuitable for decoding cognition functions whose neural features span widespread frequencies and distributed brain networks.

### 6.2. Preclinical Validation

Our ovine study validates core SNS functionality. Due to the small sheep brain, we implanted two multi-contact depth leads. Stimulation at a single contact pair demonstrated that increasing amplitude and frequency systematically alter alpha- and gamma-band activity at other recording electrodes. Personalized logistic regression classifiers reliably predicted animal movement in hold-out sessions, demonstrating in vivo that the SNS can detect and modulate behaviorally relevant neural signals. Histological analyses showed no adverse tissue response to either lead type.

### 6.3. Limitations and Future Design Considerations

Although 60 channels have proven sufficient to decode memory lapses in large datasets [24, 14], increasing channel count would likely improve classifier performance and parameter optimization. However, increasing leads beyond four would increase surgical risk, procedural complexity, and IPG size. The SNS architecture could accommodate twice the channel count, with the main constraint being connector block size.

The ovine model, while valuable for demonstrating functionality, has translation limitations. The smaller sheep brain necessitated only two shorter depth leads (with only nine bipolar pairs available for recording the effects of stimulation after excluding stimulation-adjacent contacts). Additionally, while our experiments demonstrated reliable decoding of movement and spectral power modulation, we did not exercise the SNS’s ability to decode higher-cognitive functions, such as memory or its closed-loop functionality.

### 6.4. Conclusions

The Smart Neurostimulation System (SNS) represents a major advance in neuromodulatory technology, offering simultaneous 60-channel brain activity monitoring and automatically configurable, targeted electrical stimulation based on real-time spectral analysis of neural data. Ovine data demonstrate core functionality. Although designed to evaluate an earlier proof-of-concept therapy for memory loss, the SNS’s multi-channel sensing and brain-state-contingent stimulation holds the potential to address a broad range of pathologies that exhibit moment-to-moment fluctuations, such as depression, anxiety, chronic pain, and attention disorders. The platform’s ability to record from distributed brain networks while delivering targeted stimulation to deep brain structures provides the foundation for a broad range of personalized, biomarker-guided therapies.

## Declarations

- Funding. The Army Medical Research and Development Command funded some activities under MTEC project 20-06-MOM. The views, opinions, and/or findings contained in this material are those of the authors and should not be interpreted as representing the official views or policies of the Department of Defense or the U.S. Government.
- Conflict of interest/Competing interests. D.S.R. and M.J.K. hold a greater than 5% equity interest in Nia Therapeutics Inc., a company intended to develop and commercialize brain stimulation therapies for memory restoration.

## References

[1] D. R. Nair, K. D. Laxer, P. B. Weber, A. M. Murro, Y. D. Park, G. L. Barkley, B. J. Smith, R. P. Gwinn, M. J. Doherty, K. H. Noe, R. S. Zimmerman, G. K. Bergey, W. S. Anderson, C. Heck, C. Y. Liu, R. W. Lee, T. Sadler, R. B. Duckrow, L. J. Hirsch, R. E. J. Wharen, W. Tatum, S. Srinivasan, G. M. McKhann, M. A. Agostini, A. V. Alexopoulos, B. C. Jobst, D. W. Roberts, V. Salanova, T. C. Witt, S. S. Cash, A. J. Cole, G. A. Worrell, B. N. Lundstrom, J. C. Edwards, J. J. Halford, D. C. Spencer, L. Ernst, C. T. Skidmore, M. R. Sperling, I. Miller, E. B. Geller, M. J. Berg, A. J. Fessler, P. Rutecki, A. M. Goldman, E. M. Mizrahi, R. E. Gross, D. C. Shields, T. H. Schwartz, D. R. Labar, N. B. Fountain, W. J. Elias, P. W. Olejniczak, N. R. Villemarette-Pittman, S. Eisenschenk, S. N. Roper, J. G. Boggs, T. A. Courtney, F. T. Sun, C. G. Seale, K. L. Miller, T. L. Skarpaas, M. J. Morrell, Nine-year prospective efficacy and safety of brain-responsive neurostimulation for focal epilepsy., Neurology 95 (9) (2020) e1244– e1256. doi:10.1212/WNL.0000000000010154.

[2] S. Stanslaski, R. L. S. Summers, L. Tonder, Y. Tan, M. Case, R. S. Raike, N. Morelli, T. M. Herrington, M. Beudel, J. L. Ostrem, S. Little, L. Almeida, A. Ramirez-Zamora, A. Fasano, T. Hassell, K. T. Mitchell, E. Moro, M. Gostkowski, N. Sarangmat, H. Bronte-Stewart, On behalf of the ADAPT-PD Investigators, Sensing data and methodology from the Adaptive DBS Algorithm for Personalized Therapy in Parkinson’s Disease (ADAPT-PD) clinical trial, npj Parkinson’s Disease 10 (1) (2024) 174. doi:10.1038/s41531-024-00772-5. URL https://doi.org/10.1038/s41531-024-00772-5

[3] N. A. Mekhail, R. M. Levy, T. R. Deer, L. Kapural, S. Li, K. Amirdelfan, J. E. Pope, C. W. Hunter, S. M. Rosen, S. J. Costandi, S. M. Falowski, A. H. Burgher, C. A. Gilmore, F. A. Qureshi, P. S. Staats, J. Scowcroft, T. McJunkin, J. Carlson, C. K. Kim, M. I. Yang, T. Stauss, E. A. Petersen, J. M. Hagedorn, R. Rauck, J. W. Kallewaard, G. Baranidharan, R. S. Taylor, L. Poree, D. Brounstein, R. V. Duarte, G. E. Gmel, R. Gorman, I. Gould, E. Hanson, D. M. Karantonis, A. Khurram, A. Leitner, D. Mugan, M. Obradovic, Z. Ouyang, J. Parker, P. Single, N. Soliday, Ecap-controlled closed-loop versus open-loop scs for the treatment of chronic pain: 36-month results of the evoke blinded randomized clinical trial., Reg Anesth Pain Med 49 (5) (2024) 346–354. doi:10.1136/rapm-2023-104751.

[4] H. Nijhuis, J.-W. Kallewaard, J. van de Minkelis, W.-J. Hofsté, L. Elzinga, P. Armstrong, I. Gültuna, E. Almac, G. Baranidharan, S. Nikolic, A. Gulve, J. Vesper, B. E. Dietz, D. Mugan, F. J. P. M. Huygen, Durability of evoked compound action potential (ecap)-controlled, closed-loop spinal cord stimulation (scs) in a real-world european chronic pain population., Pain Ther 13 (5) (2024) 1119–1136. doi:10.1007/s40122-024-00628-z.

[5] K. W. Scangos, A. N. Khambhati, P. M. Daly, G. S. Makhoul, L. P. Sugrue, H. Zamanian, T. X. Liu, V. R. Rao, K. K. Sellers, H. E. Dawes, P. A. Starr, A. D. Krystal, E. F. Chang, Closed-loop neuromodulation in an individual with treatment-resistant depression, Nature Medicine 27 (10) (2021) 1696–1700. doi:10.1038/s41591-021-01480-w. URL https://doi.org/10.1038/s41591-021-01480-w

[6] A. S. Widge, D. A. Malone, D. D. Dougherty, Closing the loop on deep brain stimulation for treatment-resistant depression, Frontiers in Neuroscience Volume 12–2018 (2018). doi:10.3389/fnins.2018.00175. URL https://www.frontiersin.org/journals/neuroscience/articles/10.3389/fnins.

[7] Y. Ezzyat, J. E. Kragel, J. F. Burke, D. F. Levy, A. Lyalenko, P. A. Wanda, L. O’Sullivan, K. B. Hurley, S. Busygin, I. Pedisich, M. R. Sperling, G. A. Worrell, M. T. Kucewicz, K. A. Davis, T. H. Lucas, C. S. Inman, B. C. Lega, B. C. Jobst, S. A. Sheth, K. Zaghloul, M. J. Jutras, J. M. Stein, S. R. Das, R. Gorniak, D. S. Rizzuto, M. J. Kahana, Direct brain stimulation modulates encoding states and memory performance in humans, Current Biology 27 (9) (2017) 1251–1258. doi:10.1016/j.cub.2017.03.028.

[8] Y. Ezzyat, P. A. Wanda, D. F. Levy, A. Kadel, A. Aka, I. Pedisich, M. R. Sperling, A. D. Sharan, B. C. Lega, A. Burks, R. E. Gross, C. S. Inman, B. C. Jobst, M. A. Gorenstein, K. A. Davis, G. A. Worrell, M. T. Kucewicz, J. M. Stein, R. Gorniak, S. R. Das, D. S. Rizzuto, M. J. Kahana, Closed-loop stimulation of temporal cortex rescues functional networks and improves memory, Nature Communications 9 (1) (2018) 365. doi:10.1038/s41467-017-02753-0. URL https://www.nature.com/articles/s41467-017-02753-0

[9] J. E. Kragel, Y. Ezzyat, M. R. Sperling, R. Gorniak, G. A. Worrell, B. M. Berry, C. Inman, J.-J. Lin, K. A. Davis, S. R. Das, J. M. Stein, B. C. Jobst, K. A. Zaghloul, S. A. Sheth, D. S. Rizzuto, M. J. Kahana, Similar patterns of neural activity predict memory function during encoding and retrieval, NeuroImage 155 (2017) 60–71. doi:10.1016/j.neuroimage.2017.03.042. URL https://linkinghub.elsevier.com/retrieve/pii/S1053811917302549

[10] C. T. Weidemann, J. E. Kragel, B. C. Lega, G. A. Worrell, M. R. Sperling, A. D. Sharan, B. C. Jobst, F. Khadjevand, K. A. Davis, P. A. Wanda, A. Kadel, D. S. Rizzuto, M. J. Kahana, Neural activity reveals interactions between episodic and semantic memory systems during retrieval., Journal of Experimental Psychology: General 148 (1) (2019) 1–12. doi:10.1037/xge0000480. URL https://doi.apa.org/doi/10.1037/xge0000480

[11] M. J. Kahana, Y. Ezzyat, P. A. Wanda, E. A. Solomon, R. Adamovich-Zeitlin, B. C. Lega, B. C. Jobst, R. E. Gross, K. Ding, R. R. Diaz-Arrastia, Biomarker-guided neuromodulation aids memory in traumatic brain injury, Brain Stimulation 16 (4) (2023) 1086–1093. doi:10.1016/j.brs.2023.07.002. URL https://linkinghub.elsevier.com/retrieve/pii/S1935861X23018168

[12] Y. Ezzyat, J. E. Kragel, E. A. Solomon, B. C. Lega, J. P. Aronson, B. C. Jobst, R. E. Gross, M. R. Sperling, G. A. Worrell, S. A. Sheth, P. A. Wanda, D. S. Rizzuto, M. J. Kahana, Functional and anatomical connectivity predict brain stimulation’s mnemonic effects, Cerebral Cortex 34 (1) (2023) bhad427. doi:10.1093/cercor/bhad427. URL https://academic.oup.com/cercor/article/doi/10.1093/cercor/bhad427/745734

[13] M. J. Kahana, E. V. Aggarwal, T. D. Phan, The variability puzzle in human memory, Journal of Experimental Psychology: Learning, Memory, and Cognition 44 (12) (2018) 1857–1863. https://doi.org/10.1037/xlm0000553.

[14] D. Y. Rubinstein, C. T. Weidemann, M. R. Sperling, M. J. Kahana, Direct brain recordings suggest a causal subsequent-memory effect, Cerebral Cortex 33 (11) (2023) 6891–6901. doi:10.1093/cercor/bhad008. URL https://academic.oup.com/cercor/article/33/11/6891/7005169

[15] C. T. Weidemann, M. J. Kahana, Neural measures of subsequent memory reflect endogenous variability in cognitive function., Journal of Experimental Psychology: Learning, Memory, and Cognition 47 (4) (2021) 641–651. doi:10.1037/xlm0000966.

16. [16] Y. Ezzyat, N. Suthana, Brain Stimulation, in: M. J. Kahana, A. D. Wagner (Eds.), Oxford Handbook of Human Memory, 2nd Edition, Oxford University Press, 2022.

[17] J. F. Burke, A. G. Ramayya, M. J. Kahana, Human intracranial high-frequency activity during memory processing: neural oscillations or stochastic volatility?, Current Opinion in Neurobiology 31 (2015) 104–110. doi:10.1016/j.conb.2014.09.003. URL https://linkinghub.elsevier.com/retrieve/pii/S0959438814001810

[18] J. A. Greenberg, J. F. Burke, R. Haque, M. J. Kahana, K. A. Zaghloul, Decreases in theta and increases in high frequency activity underlie associative memory encoding., NeuroImage 114 (2015) 257–263. doi:10.1016/j.neuroimage.2015.03.077.

[19] U. R. Mohan, A. J. Watrous, J. F. Miller, B. C. Lega, M. R. Sperling, G. A. Worrell, R. E. Gross, K. A. Zaghloul, B. C. Jobst, K. A. Davis, S. A. Sheth, J. M. Stein, S. R. Das, R. Gorniak, P. A. Wanda, D. S. Rizzuto, M. J. Kahana, J. Jacobs, The effects of direct brain stimulation in humans depend on frequency, amplitude, and white-matter proximity, Brain Stimulation 13 (5) (2020) 1183–1195, number: 5 MohaEtal20. doi:10.1016/j.brs.2020.05.009. URL https://linkinghub.elsevier.com/retrieve/pii/S1935861X2030108X

[20] B. C. Johnson, S. Gambini, I. Izyumin, A. Moin, A. Zhou, G. Alexandrov, S. R. Santacruz, J. M. Rabaey, J. M. Carmena, R. Muller, An implantable 700µW 64-channel neuromodulation IC for simultaneous recording and stimulation with rapid artifact recovery, in: 2017 Symposium on VLSI Circuits, 2017, pp. C48–C49. doi:10.23919/VLSIC.2017.8008543. URL https://ieeexplore.ieee.org/document/8008543

[21] A. Zhou, S. R. Santacruz, B. C. Johnson, G. Alexandrov, A. Moin, F. L. Burghardt, J. M. Rabaey, J. M. Carmena, R. Muller, A wireless and artefact-free 128-channel neuromodulation device for closed-loop stimulation and recording in non-human primates, Nature Biomedical Engineering 3 (1) (2019) 15–26. doi:10.1038/s41551-018-0323-x. URL https://www.nature.com/articles/s41551-018-0323-x

[22] A. Ella, M. Keller, Construction of an MRI 3D high resolution sheep brain template, Magnetic Resonance Imaging 33 (10) (2015) 1329–1337. doi:10.1016/j.mri.2015.09.001.

[23] A. Ella, J. A. Delgadillo, P. Chemineau, M. Keller, Computation of a high-resolution MRI 3D stereotaxic atlas of the sheep brain, The Journal of Comparative Neurology 525 (3) (2017) 676–692. doi:10.1002/cne.24079.

[24] J. H. Rudoler, N. A. Herweg, M. J. Kahana, Hippocampal Theta and Episodic Memory, The Journal of Neuroscience 43 (4) (2023) 613–620. doi:10.1523/JNEUROSCI.1045-22.2022. URL https://www.jneurosci.org/lookup/doi/10.1523/JNEUROSCI.1045-22.2022

[25] S. Haufe, F. Meinecke, K. Görgen, S. Dähne, J.-D. Haynes, B. Blankertz, F. Bießmann, On the interpretation of weight vectors of linear models in multivariate neuroimaging, NeuroImage 87 (2014) 96–110. doi:10.1016/j.neuroimage.2013.10.067. URL https://linkinghub.elsevier.com/retrieve/pii/S1053811913010914

[26] N. M. Long, M. J. Kahana, Modulation of task demands suggests that semantic processing interferes with the formation of episodic associations, Journal of Experimental Psychology. Learning, Memory, and Cognition 43 (2) (2017) 167–176. doi:10.1037/xlm0000300.

[27] J. R. Manning, J. Jacobs, I. Fried, M. J. Kahana, Broadband shifts in local field potential power spectra are correlated with single-neuron spiking in humans, Journal of Neuroscience 29 (43) (2009) 13613–13620. doi:10.1523/JNEUROSCI.2041-09.2009.

[28] Y. Benjamini, D. Yekutieli, The control of the false discovery rate in multiple testing under dependency, Annals of statistics (2001) 1165–1188 Publisher: JSTOR.

[29] J. A. Roa, L. Marcuse, M. Fields, M. L. Vega-Talbott, J. Y. Yoo, S. M. Wolf, P. McGoldrick, S. Ghatan, F. Panov, Long-term outcomes after responsive neurostimulation for treatment of refractory epilepsy: a single-center experience of 100 cases., J Neurosurg 139 (5) (2023) 1463–1470. doi:10.3171/2023.2.JNS222116.

[30] S. W. Stull, J. Mogle, J. W. Bertz, A. J. Burgess-Hull, L. V. Panlilio, S. T. Lanza, K. L. Preston, D. H. Epstein, Variability in intensively assessed mood: Systematic sources and factor structure in outpatients with opioid use disorder, Psychological Assessment 34 (10) (2022) 966–977, number: 10 StulEtal22. doi:10.1037/pas0001160.

[31] J. J. Sakon, D. J. Halpern, D. R. Schonhaut, M. J. Kahana, Human hippocampal ripples signal encoding of episodic memories, The Journal of Neuroscience (2024) e0111232023doi:10.1523/JNEUROSCI.0111-23.2023. URL https://www.jneurosci.org/lookup/doi/10.1523/JNEUROSCI.0111-23.2023

[32] V. Rahimzadeh, K. M. Jones, M. A. Majumder, M. J. Kahana, U. Rutishauser, Z. M. Williams, S. S. Cash, A. C. Paulk, J. Zheng, M. S. Beauchamp, J. L. Collinger, N. Pouratian, A. L. McGuire, S. A. Sheth, R. Adolphs, R. A. Andersen, G. Baltuch, P. Brunner, S. S. Cash, E. Chang, J. L. Collinger, N. Crone, E. Fedorenko, I. Fried, J. Gold, J. Henderson, L. Hochberg, M. Howard, M. J. Kahana, J. Magnotti, A. Mamelak, N. Pouratian, R. M. Richardson, U. Rutishauser, G. Schalk, C. Schroeder, K. Shenoy, S. A. Sheth, N. Suthana, N. Tandon, Z. M. Williams, J. Wolpaw, Benefits of sharing neurophysiology data from the BRAIN Initiative Research Opportunities in Humans Consortium, Neuron 111 (23) (2023) 3710–3715. doi:10.1016/j.neuron.2023.09.029. URL https://linkinghub.elsevier.com/retrieve/pii/S0896627323007171

[33] Z. T. Sanger, T. R. Henry, M. C. Park, D. Darrow, R. A. McGovern, T. I. Netoff, Neural signal data collection and analysis of Percept™ PC Brain-Sense recordings for thalamic stimulation in epilepsy, Journal of Neural Engineering 21 (1) (2024) 012001. doi:10.1088/1741-2552/ad1dc3. URL https://dx.doi.org/10.1088/1741-2552/ad1dc3

